# Amaranth: Enhanced Single-Cell Transcript Assembly via Discriminative Modeling of UMI Reads and Internal Reads

**DOI:** 10.1101/2025.11.24.690228

**Authors:** Xiaofei Carl Zang, Tasfia Zahin, Irtesam Mahmud Khan, Qian Shi, Yi Xing, Mingfu Shao

## Abstract

**Motivation:** Single-cell RNA sequencing has transformed transcriptome profiling at cellular resolution, yet accurate reconstruction of full-length transcripts for individual cells remains a central challenge. Emerging scRNA-seq protocols can produce reads that span entire transcripts, enabling isoform-level expression analysis. For example, Smart-seq protocols combine UMI-linked reads that index and stitch together multiple reads from the same molecule, with internal reads filling coverage gaps. We demonstrate that these read types exhibit markedly different biological and statistical properties in strandness, 5’/3’ coverage bias, and genomic locality. Existing assemblers fail to leverage these distinctions, yielding suboptimal assembly.

**Results:** We developed Amaranth, a novel single-cell assembler that discriminatively models UMI and internal reads. Amaranth implements heuristics specifically designed to address the distinct biases of UMI-linked and internal reads, enabling accurate strandness assignment for internal reads, reliable splicing graph refinement, and precise transcript start site determination. We also developed Amaranth-meta, which integrates information across cells to enhance individual cell assemblies. Benchmarked on Smart-seq3 datasets from human HEK293T and mouse fibroblast cells, Amaranth outperformed other state-of-the-art assemblers in assembling individual cells and in meta-assembly. Amaranth advances isoform-level analysis in single-cell transcriptomics, facilitating detailed studies at cellular resolution.

**Availability and Implementation:** Amaranth is implemented in C++ and is freely available at https://github.com/Shao-Group/amaranth under the BSD-3-Clause license. Scripts, documentation, and data for reproducing experiments in this manuscript are available at https://github.com/Shao-Group/amaranth-test.

## 1 Introduction

Single-cell RNA-seq (scRNA-seq) technologies have been widely adopted for transcriptomic profiling at single-cell resolution, enabling the study of RNA velocity [15], cellular heterogeneity [37], spatial analysis [35, 24], and cell-cell communication [6] with unprecedented detail. Novel scRNA-seq protocols are rapidly being developed in both academia and industry. Droplet-based microfluidic methods are among the dominant approaches for scRNA-seq [4], e.g., 10x Genomics Chromium platform [44], Drop-seq [19], and SPLiT-seq [25]. Although high-throughput and cost-effective, those methods are often biased to either the 3’-end or the 5’-end of the RNA molecules. Most droplet-based protocols capture only one end of each RNA molecule because cell barcodes and unique molecular identifiers (UMIs) are ligated or reverse-transcribed at one terminus, leaving the internal portions of transcripts sparsely covered. Consequently, these methods are largely limited to gene-level analysis because the read coverage at splice junctions, which is essential for distinguishing isoforms originating from the same gene, is typically very low.

Nevertheless, there is great interest in pinpointing isoform alternative splicing and quantification in single cells at a finer resolution. To meet this goal, methods that generate reads covering entire transcripts with reduced bias are increasingly available. These include short-read approaches such as RamDA-seq [11], the Smart-seq series [23, 9, 8], FLASH-seq [10], and VASA-seq [27], as well as long-read protocols including ScISOr-seq [7] and HIT-scISOseq [32]. These protocols and their alternatives are becoming increasingly widespread and have been used to study transcript architecture and splice variants [39]. Most of them rely heavily on UMI-based techniques. For example, in the Smart-seq3 protocol, two types of reads, referred to as UMI reads and internal reads, are produced. UMI reads are linked to UMIs that index and link multiple reads from the same unique molecule. Similar to droplet-based protocols, Smart-seq3 UMI reads capture only the 5’-end of RNAs. On the other hand, internal reads resemble bulk RNA-seq data, providing increased and even coverage in the internal part and the 3’-end of RNA molecules, despite containing more noise [9]. By combining both types of reads, we are given a unique opportunity to achieve isoform-level resolution in single-cell RNA-seq analysis.

The reconstruction of full-length transcripts from RNA-seq reads is a well-known problem — transcript assembly. Whilst numerous transcript assembly tools have been developed in the past decades, they have overlooked the specialty embedded in scRNA-seq data. Tools designed for bulk paired-end RNA-seq data (e.g., StringTie2 [14], CLASS2 [33], cufflinks [36], TransComb [17], to just name a few) or multi-end reads (e.g., Scallop2 [43]) pool all reads together regardless of the difference between UMI and internal reads. Bulk RNA-seq assemblers are also unaware of the sparse coverage and dropout events in scRNA-seq, producing partial and fragmented transcripts. Assemblers specific to scRNAseq data (e.g., scRNAss [18] and RNA-Bloom [20]) often rely on the reference transcriptome annotation to fill the coverage gaps, but this process is exposed to reference bias and *ad hoc* assumptions. Meta-assemblers (e.g., Aletsch [30], TransMeta [41], PsiCLASS [34]) aim to assemble transcripts from multiple RNA-seq samples. They can cross-reference information from other cells, but they tend to reconstruct consensus transcripts and also ignore the differences between UMI and internal scRNA-seq reads. Beaver [31] is a recent method that is specifically designed to assemble scRNA-seq transcripts without relying on reference annotation, but it does not directly assemble reads or alignments. Instead, Beaver takes single-cell transcriptomes from either a single-sample or a meta-assemblers and extrapolates cross-cell information to augment them. It is dependent on and benefits from a high-quality upstream assembly.

In this work, we contribute to the scRNA-seq assembly problem in two aspects. First, our work highlights that the two types of scRNA-seq reads, UMI or internal, display distinct biological and statistical properties. While UMI reads exhibit distribution biases towards 5’-end and coverage sparsity, they are highly specific and precise. In contrast, internal reads resemble bulk RNA-seq data, with an increased sequencing coverage and a higher proportion of reads from unspliced introns. It is of keen interest to separate those two types of reads, and to model them differently. Second, we introduced Amaranth, a novel assembler that addresses the critical gap of scRNA-seq assembly. Amaranth discriminatively integrates UMI and internal reads to achieve accurate, cellspecific transcript assembly without relying on external references. Amaranth can perform transcript assembly on individual cell basis (without other cells) or perform a meta-assembly using all cells’ information. We show that on human and mouse Smart-seq3 datasets, Amaranth outperforms the state-of-the-art tools, including Scallop2, StringTie2, Aletsch, and PsiCLASS, in both single-cell assemblies and meta-assembly.

## 2 Disparities between UMI reads and internal reads

We first investigated the statistical and biological disparities between UMI and internal reads. These new observations motivated and provided insights for our new assembler Amaranth, described in Section 3.

The data we analyzed were two Smart-seq3 scRNA-seq datasets consisting of 192 human HEK293T cells and 369 mouse fibroblast cells respectively [9]. In both datasets, there are more internal reads than UMI reads in total reads. The great majority of the cells have more internal reads in both human and mouse datasets. On average, 75% of reads in a human HEK293T cell and 71% of reads in a mouse fibroblast cell are internal, while only 25% of reads in a human cell and 29% of reads in a mouse cell are UMI reads (see Fig. S1).

We first show that, UMI reads are strand-specific whereas internal reads are not. Specifically, we used RSeQC [38] to determine the reads’ strandness (`infer_experiment.py`). The strandness of more than 92% reads can be successfully determined in both human and mouse Smart-seq3 datasets. The UMI reads are highly specific in FR-strandness. On average, 82%–89% of UMI reads are FR-stranded and only 5%–11% of UMI reads are RF-stranded in human and mouse datasets. Conversely, the internal reads lose their strand specificity, as they roughly have the same amount of RF-stranded reads and FR-stranded reads. Those observations are present in almost all cells in both datasets (see Fig. 1A-B and Fig. S2A-B).

**Figure 1.**
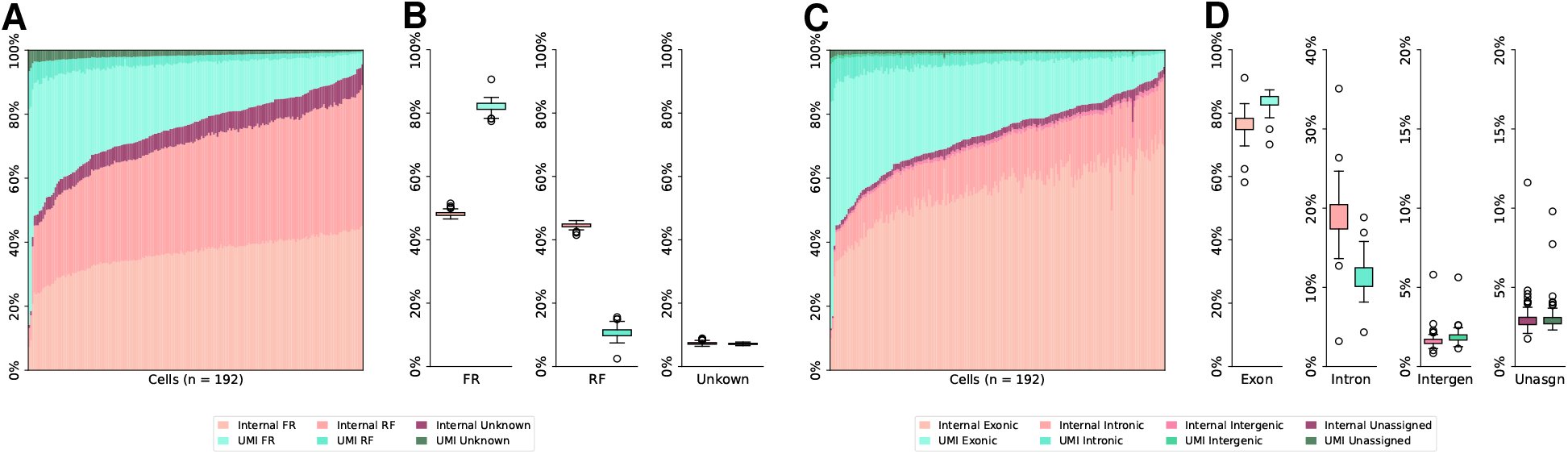
Read strandness and genome feature distributions of UMI and internal reads in human HEK293T cells. **(A)** Strandness of reads in each HEK293T cell. Cells are sorted in ascending order of the internal reads proportion. **(B)** Breakdown of read strandness by read type. Internal reads: 48.16% FR-strandness, 44.50% RF-strandness, 7.34% unknown. UMI reads: 82.13% FR-strandness, 10.73% RF-strandness, 7.14% unknown. **(C)** Genome features of read tags in each HEK293T cell. Cells are sorted in ascending order of the internal reads proportion. **(D)** Breakdown of genome feature distributions by read type in HEK293T cells. Internal read tags: 76.42% exonic, 19.01% intronic, 1.62% intergenic, 2.95% unassigned. UMI read tags: 83.75% exonic, 11.40% intronic, 1.87% intergenic, 2.98% unassigned.

Next, we examine the locality of UMI reads and internal reads. A read tag will be assigned to one genome feature, with precedence given to exonic regions, intronic regions, intergenic regions within 10 kb, or unassigned regions, inferred using RSeQC (`read_distribution.py`). The main reason that a read tag cannot be assigned is that it is intergenic and beyond a 10kb window from any gene or within a 10kb window between two genes. Both human HEK293T and mouse fibroblast datasets are evaluated and illustrated respectively in Fig. 1C-D and Fig. S2C-D. The genome feature distributions of UMI and internal reads are different, particularly in the HEK293T dataset (Fig. 1C- D). Quantitatively, the proportion of internal read tags mapping to intronic regions is almost twice that of UMI read tags, approximately 19% and 11% in the HEK293T Smart-seq3 dataset (Fig. 1D). Considering the internal reads are 3 times as abundant as UMI reads, approximately 84% of intronic contamination is contributed by internal reads in the human dataset. This trend is not as evident in the mouse fibroblast dataset, except for a lower variance of the ratios of intronic UMI read tags. This is possibly because the mouse transcriptome is relatively simpler with fewer and shorter introns [1, 2, 26].

An example of read alignments and distributions is shown in Fig. 2. UMI reads demonstrate a clear 5’-end bias and strand specificity as expected, whilst internal reads are relatively evenly distributed along the gene body but include a higher proportion of intronic alignments. These observations are consistent with our analysis and underscore the necessity of a discriminative model to leverage the strengths of both read types while mitigating the impact of internal read contamination.

**Figure 2.**
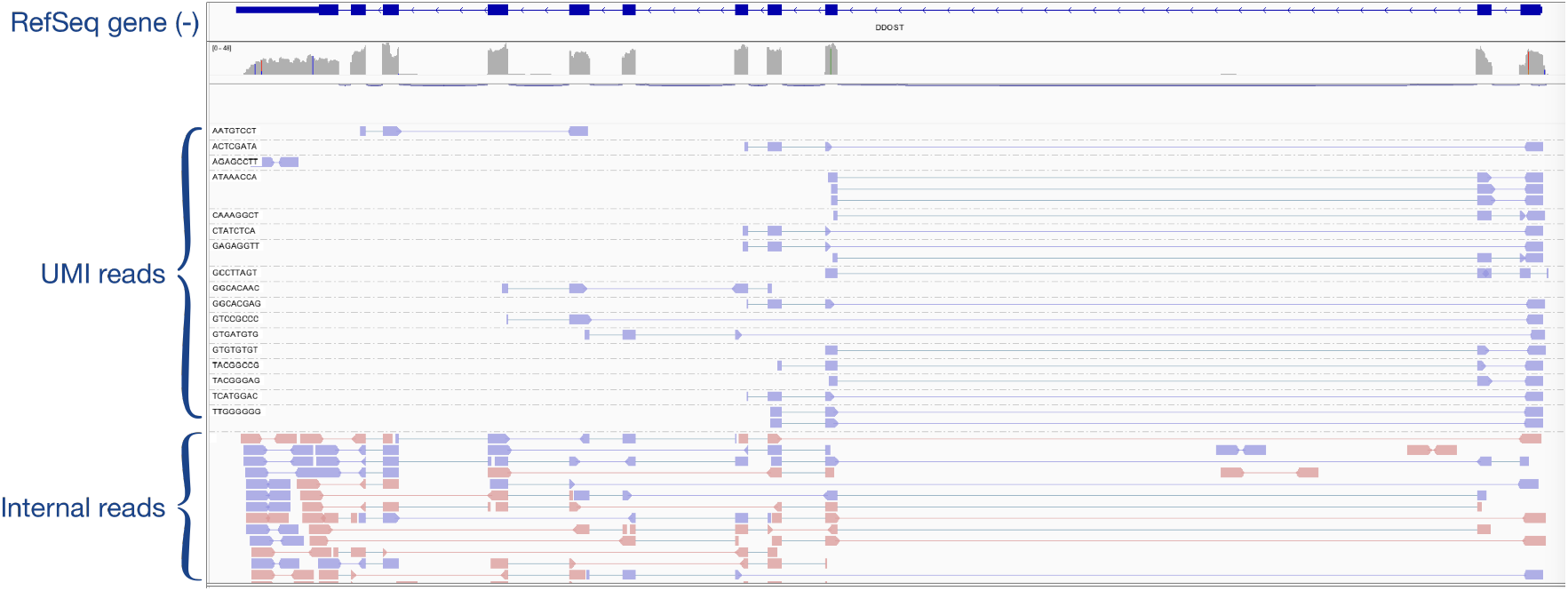
Example screenshot of Smart-seq3 read alignments using IGV. The gene is annotated in the RefSeq GRCh38 transcriptome annotation and coded in the negative strand of the genome. Violet read pairs are FR-stranded, whilst pink read pairs are RF-stranded. UMI reads exhibit a 5’ bias, while internal reads provide relatively even coverage but include noise. UMI reads are FR-strand-specific, while internal reads are not strand-specific.

## 3 Amaranth

Amaranth is a reference-based assembler that takes demultiplexed single-cell RNA-seq read alignments to a reference genome (*i*.*e*. a *bam* file) and outputs a set of assembled transcripts (*i*.*e*. a *gtf* file) for each individual cell. Amaranth follows an algorithmic framework that first builds the splice graph from read alignment and subsequently performs graph-decomposition, an approach that is also employed in other assemblers, including the Scallop series [28, 43]. The novel contribution of Amaranth is that, beyond the assembly of all RNA-seq reads, it employs a multi-tiered computational procedure to model the disparities in the UMI-linked and internal reads. It first starts with grouping overlapping and proximate reads into gene loci and correcting the strandness of ambiguous internal reads with respect to nearby UMI reads, and filters PCR duplicates and potential contaminating loci. Second, a splice graph is constructed using both UMI and internal reads, followed by the purge of potential intronic contamination. Third, precise detection of transcription start sites leverages UMI read termini to anchor the isoform boundary.

### 3.1 Reads classifications and corrections

Amaranth first distinguishes UMI reads based on the `UB` optional field tag in the *bam* file, which contains their UMI sequences. UMI reads from the same original molecule should have the same UMI sequence, namely, the same `UB` value. Since the UMI information of internal reads is lost during the fragmentation procedure, internal reads do not have `UB` tag or have empty `UB` value.

UMI reads are known to be strand-specific [9] and can be easily determined. However, internal reads present a manner of “bulk-like” features and lose the strand-specificity (see details in Section 2). Some internal reads contain splicing junction(s), in which case their strandness can be inferred from the intron motif by splice-aware aligners, such as STAR [5], HISAT [12, 13], and minimap2 [16]. For internal reads that do not contain splicing information to deduce strandness, Amaranth infers their strandness by leveraging the strandness of nearby UMI reads. Specifically, strandness of genes is determined by the gene’s UMI reads and strand-resolved internal reads (e.g., using an intron motif). Then, Amaranth assigns strand-unknown internal reads to nearby gene(s) within 100 bp and determines the strandness accordingly. In rare cases, if there exist multiple nearby genes from both strands, the aforementioned internal reads will be assigned to both strands and both genes.

To reduce noise, Amaranth introduced a new procedure to remove fragment duplicates that are caused by PCR-based sequencing methods. Putative PCR duplicates are determined if two read pairs have exactly the same alignment to the reference genome, *i*.*e*., identical position and CIGAR string. Duplicates are also removed if marked by the aligner or an optional processing tool, such as picard [3]. After read classifications and corrections, UMI reads are expected to satisfactorily support a gene in the sense of read count and ratio. If a gene locus has too few UMI reads and/or too low a proportion of UMI reads, such gene loci are likely intergenic or intronic contamination by internal reads and will be purged.

### 3.2 Splice graph construction and contamination removal

Amaranth uses all reads in a gene locus, including both UMI and internal reads, to construct the splice graph, a data structure that is essential for transcript assembly. A splice graph is a weighted, directed acyclic graph *G* = (*V, E*) and associated weights *w*_*v*_, *w*_*e*_, and *w*_*m*_. Each aligned region will be presumed as a partial or full exon, represented as a vertex *v* in *V* . The average read coverage in this region is represented as the node weight of *v*, denoted as *w*_*v*_(*v*). If two vertices *u* and *v* are connected by a read via splicing or reading through, we add one directed edge *e* = (*u, v*) to *E*, assuming the coordinates of *u* precede those of *v*. The quantity of reads connecting *u* and *v* is represented as edge weight, denoted as *w*_*e*_(*e*). The average UMI read coverage of vertex *v* is represented as vertex UMI support, denoted as *w*_*m*_(*v*). We add a source *s* and sink *t* to *V* and connect *s*/*t* to possible starting/ending vertices, respectively.

We showed that internal reads introduce more noise and contamination from intronic regions. Such intronic noise may result in false-positive transcripts with intron retention, which is one of the major sources of transcript assembly errors. Amaranth hence introduces a rule-based procedure to prune putative intronic artifacts (previously considered partial exons) to circumvent contaminations. This procedure is similar to intron retention removal defined in Ref. [42], which filters post-assembly full-length transcripts. Nevertheless, Amaranth prunes contamination prior to assembly and utilizes the proximate information of vertices (see below), that will be lost post-assembly. This prior-to-assembly pruning reduces graph complexity and improves assembly in both accuracy and resource consumption. The intron retention removal is formally defined as follows.

Given two vertices *u, v* ∈ *V* in the splice graph *G*, denote the genomic regions that they respectively span as left-close, right-open intervals, [*u*_*l*_, *u*_*r*_), and [*v*_*l*_, *v*_*r*_). Assuming *u* topologically precedes *v* in *G*, we say that *u* is immediately joint to *v* if *u*_*r*_ = *v*_*l*_. That means the partial exons represented by *u* and *v* are consecutive regions in the genome. A vertex *v* ∈ *V* will be considered as a candidate for *full* intron retention if the following four conditions are met. (1.) *v* has exactly one incoming edge and one outgoing edge. Denote the preceding and succeeding neighbors of *v* respectively as *v*^−^ and *v*^+^. (2.) *v* is not adjacent to source *s* nor sink *t* (i.e., *v* represents an internal putative partial exon). (3.) *v* is immediately joint to both *v*^−^ and *v*^+^. (4.) *v* is skipped by a direct edge between its two neighbors, meaning an edge *e*^*′*^ = (*v*^−^, *v*^+^) ∈ *E* exists (e.g. *v*_3_ in Fig. 3B). Hence, *v* potentially represents a full retention of the intron 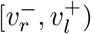 instead of a real partial exon. Intuitively, if *v* is an artifact from an unspliced intron, its vertex weight and edge weights should be lower than a threshold times the weight of the skipping edge. We consider *v* as a retained intron and remove it from the splice graph when it is skipped by a high weight edge: *w*_*v*_(*v*) ≤ *a*_1_ × *w*_*e*_(*v*^−^, *v*^+^) or *w*_*e*_(*v*^−^, *v*) ≤ *a*_2_ × *w*_*e*_(*v*^−^, *v*^+^) or *w*_*e*_(*v, v*^+^) ≤ *a*_3_ × *w*_*e*_(*v*^−^, *v*^+^); and lowly supported by UMI reads: *w*_*m*_(*v*) ≤ *a*_4_. Providing the possibility that *v* is from contamination by PCR amplification of an intron lariat, *v*’s node weight will be significantly greater than its edge weights. Thus, we also consider that *v* is a retained intron when *w*_*v*_(*v*) ≥ *a*_5_ × max{*w*_*e*_(*v*^−^, *v*), *w*_*e*_(*v, v*^+^)}. (*a*_1_, *a*_2_, · · ·, *a*_5_) are user-defined parameters whose default values are available in Note S1.

**Figure 3.**
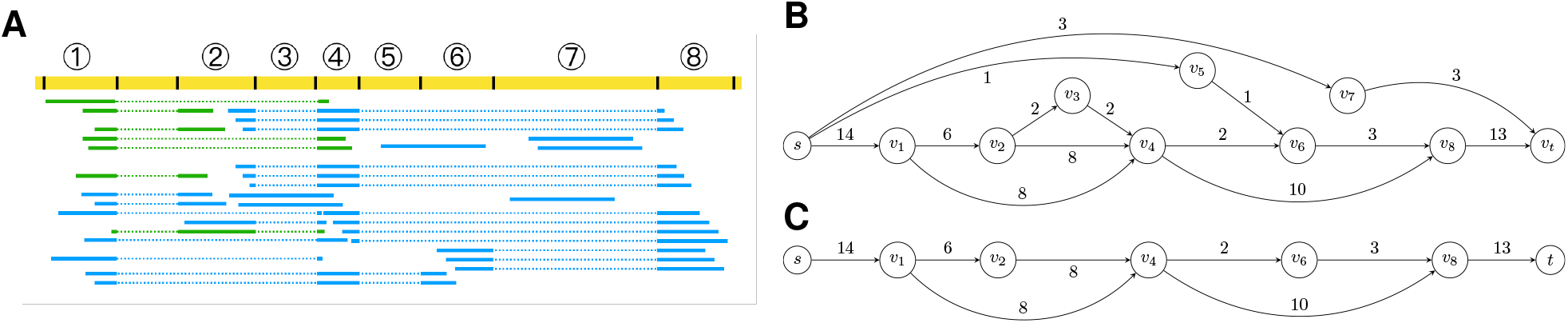
Splice graph construction and pruning. **(A)** UMI reads (green) and internal reads (blue) are aligned to a locus in the genome (yellow). Some reads are spliced into multiple tags. Aligned read tags are represented as solid lines, and splice junctions are represented as dotted lines. Inferred splice sites are marked with black bars. Genomic regions are split at splice sites into non-overlapping regions. Subsequently, those regions aligned by read tag(s) are considered putative (partial) exons and marked with serialized indices (1, 2,…, 8). **(B)** Constructed splice graph for the read alignments. Putative exons are represented as nodes with their indices marked inside each node. Edge weights marked near each edge indicate the number of reads supporting a splice junction. Node weight *w*_*v*_(·) and UMI support *w*_*m*_(·) of each node are not shown in this figure. **(C)** Splice graph after intron retention removal. Vertex *v*_3_ is removed as full intron retention. Vertex *v*_5_ is removed as partial intron retention. Vertex *v*_7_ is removed as intron contamination. Note that all of *v*_1_, *v*_5_ and *v*_7_ introduce potential alternative first exons, represented as an in-edge from *s*, but only *v*_1_ is a true first exon that is well supported by UMI reads.

Sometimes an unspliced intron is not fully covered by sequencing reads. Since the middle part of the intron is absent, a *partial* intron retention might appear as a false-positive alternative first exon, or, respectively, a false-positive alternative last exon (e.g. *v*_5_ in Fig. 3B). For simplicity and illustration purposes, we list only the criteria of the former situation, as the latter situation is symmetric. A vertex *v* will be considered as a candidate for partial intron retention if it meets the aforementioned condition (1.), and three new conditions (2.) *v* is adjacent to source *s*; (3.) *v* is immediately joint to its other succeeding neighbor *v*^+^; and (4.) *v*^+^ has at least 2 in-edges (i.e. *v*^+^ has in-edges other than from *v*). Denote the maximal weight of the in-edges of *v*^+^ as *w*^*′*^, i.e. *w*^*′*^ = max{*w*_*e*_(*u, v*^+^) | (*u, v*^+^) ∈ *E*}. Similar to a full intron retention, a partially retained intron should have lower node weight and edge weights than *w*^*′*^ and be lowly supported by UMI reads. That means *w*_*v*_(*v*) ≤ *b*_1_ × *w*^*′*^ or *w*_*e*_(*v, v*^+^) ≤ *b*_2_ × *w*^*′*^, and *w*_*m*_(*v*) ≤ *b*_3_, where *b*_1_ *b*_2_, and *b*_3_ are user-defined parameters (see also in Note S1).

Note that single vertices that are adjacent to both *s* and *t* are not considered for full or partial intron retentions in the previous steps (e.g. *v*_7_ in Fig. 3B). They are likely a result of a single-exon transcript or contamination from an intron lariat. In either case, it is easy to isolate such vertices from the splice graph, since they are not connected. A true single-exon transcript should be well supported with a high UMI-read support value *w*_*m*_(·).

### 3.3 First exon identification and transcript selection

The refined splice graph from the previous step will be decomposed into paths in a phase-preserving manner as described in Ref. [28]. Those paths are candidate full-length transcripts. Amaranth introduces a filtering step using UMI reads. It has been known that UMI scRNA-seq reads are very biased towards the end of a transcript, as UMI sequences are appended to one end of the RNA molecules during library preparation [44, 9]. Hence, UMI reads can accurately indicate the transcript start or end site, depending on which end it is biased towards. In the specific case of Smart-seq3, the UMIs are attached to the 5’ end of an RNA, so UMI reads indicate transcript start sites, *i*.*e*. position of the first exon. The candidate transcripts whose first exon is not supported by UMI reads are likely artifacts arising from intergenic or intronic contamination. This type of contamination is primarily caused by the nonselective and unspecific sequencing of internal reads. Amaranth anchors transcript start sites by requiring assembled transcripts to have their first exon aligned and supported by a certain number of UMI reads (tunable parameter, default: 1). At the end, Amaranth collects all paths that passed all filtering criteria and considers them full-length transcripts.

### 3.4 Amaranth and Amaranth-meta

Amaranth can be run as a single-cell assembler to assemble one single cell each time independent of other cells. We also extend Amaranth as a meta-assembler, referred to as Amaranth-meta, that takes all cells as input and generates an enhanced assembly for individual cells. In Amaranth-meta, all reads are first pooled as a “super-cell”, followed by assembling the expressed transcripts. Amaranth-meta assigns a transcript to a cell if a substantial number of exons is supported by reads from this specific cell (default: 30%, tunable parameter). The assigned transcripts are then unioned with transcripts assembled on individual cells (using the single-cell mode of Amaranth) to obtain the enhanced assembly for the cell. Fig. S3 illustrates the outlines of Amaranth algorithm.

## 4 Results

### 4.1 Datasets and Pipeline

We compare the performance of Amaranth with other methods using a pipeline illustrated in Fig. 4. The evaluation was conducted on the same two Smart-seq3 datasets [9]. The two datasets were pre-processed in a way similar to that previously described in Ref. [9]. Briefly, reads were demultiplexed using zUMIs [21] into individual cells, and then the reads of each cell were aligned to the human reference genome GRCh38 or mouse reference genome GRCm39 using STAR [5]. Cell barcodes of each read are stored in the `BC` tag in the *bam* file. Reads with the same `BC` tags are from the same cell. UMI sequences for applicable reads are stored in the `UB` tag in the *bam* file. Reads with the same `UB` tags are UMI reads and are from the same unique RNA molecule. Reads without valid `UB` tags are considered internal reads. The read alignment *bam* file is split into individual files according to their `BC` tags. Each of these split *bam* files consists of read alignments from the same cell.

**Figure 4.**
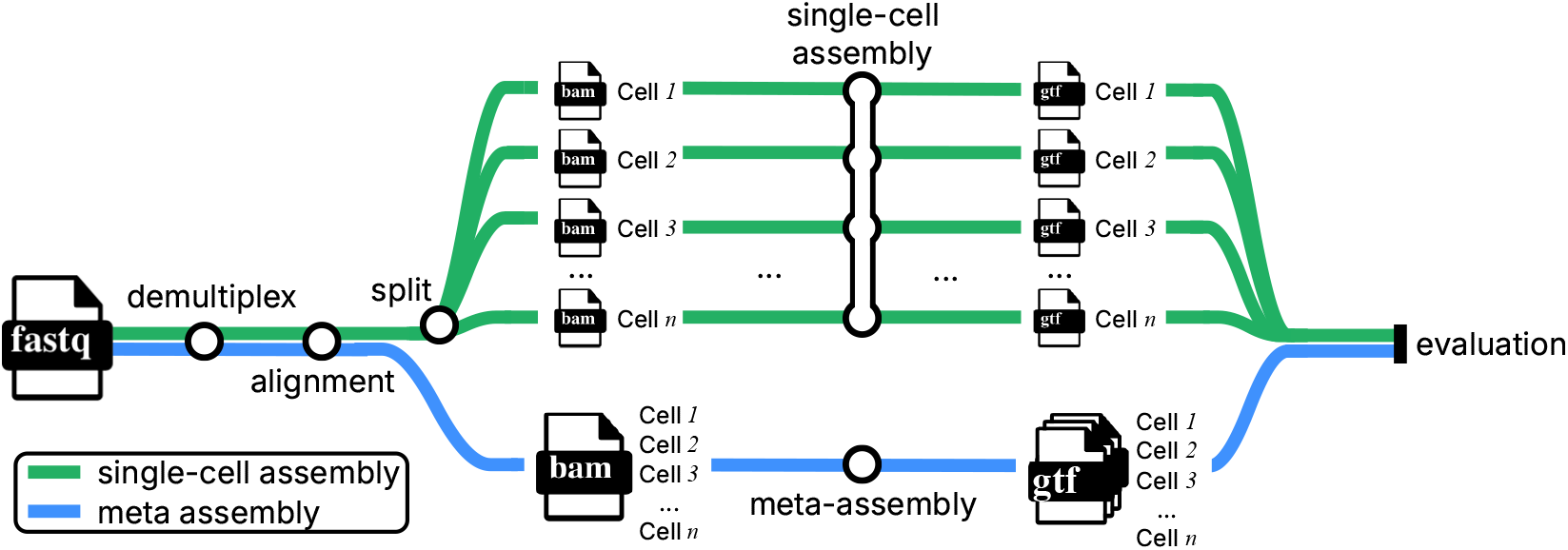
Overview of experiment workflows. Smart-seq3 scRNA-Seq reads are demultiplexed and then aligned to the reference genome using zUMIs [21] and STAR [5]. In the single-cell assembly workflow, read alignments are split into individual *bam* files. Each of these *bam* files consists of all alignments from a unique single cell. Assembly tools such as Amaranth take each *bam* file as input and output assembled transcripts for this specific cell in *gtf* format. All cells are assembled separately. In the meta-assembly workflow, meta-assemblers extrapolate other cells’ alignments to improve the assembly. All cells’ alignments are collected together and piped to downstream meta-assemblers, e.g., Amaranth-meta. At the end of the workflow, all assembled transcripts are evaluated against the reference transcriptome using gffcompare.

All assembly tools take one cell’s alignment *bam* files as input and were run with default parameters to assemble transcripts (*i*.*e*. outputting *gtf* files) for each cell in both human HEK293T and mouse fibroblast datasets. All meta-assembly tools take all cells’ alignment files at once and assemble transcripts for all cells. Gffcompare [22] was used to evaluate the assembled transcripts against reference transcriptome annotations (RefSeq GRCh38 for human or GRCm39 for mouse). Gffcompare classifies a multi-exon transcript as “matching” if its intron chain is identical to that of an annotated transcript. “Transcript level precision” (the proportion of assembled transcripts that match the reference) and “number of matching transcripts” (proportional to sensitivity) were reported. Note that the majority of annotated transcripts are not expressed and/or not sequenced in one specific cell. The traditional sensitivity, defined as the proportion of matching transcripts to all reference transcripts, will be inherently very small and uninformative for scRNA-seq, regardless of algorithms. Instead, we report the absolute count of matching transcripts, which directly measures how many true isoforms each tool successfully reconstructs.

To account for the inherent trade-offs between sensitivity and precision across different assembly methods, we compared adjusted precisions under controlled sensitivities. This is an established approach and is necessary to benchmark the performance of different tools [42]. We employed a coverage-based filtering strategy, which has been previously used and recognized as an effective way of adjusting precisions [28, 42, 30]. This approach is based on the principle that transcripts with higher predicted coverage are more likely to represent true transcripts. For each cell, we identified the method with the smallest number of matching transcripts (i.e. lowest recall) and used this number as a baseline. For each of the other methods, assembled transcripts were sequentially removed in ascending order of coverage until their sensitivity matched this baseline, thereby enabling fair precision comparisons at equivalent sensitivity levels.

### 4.2 Amaranth improves transcript assembly of individual cells

We compare Amaranth with two popular assemblers, StringTie2 [14] and Scallop2 [43], on individual cells. The assembly accuracy across all cells on both datasets is illustrated in Fig. 5. Compared to Scallop2, Amaranth highly increased assembly precision by approximately 10%-15% on the majority of those 192 HEK293T cells and 369 mouse fibroblast cells, illustrated by the higher y-axis values (precision) of scattered dots in Fig. 5A and Fig. 5E. Amaranth and Scallop2 have much higher precision than StringTie2. Under the same controlled sensitivity, Amaranth leads the precision in nearly all of those individual cells in both datasets (Fig. 5B,F). Amaranth’s average controlled precision is 72.97%, while Scallop2 and StringTie2 achieved average controlled precisions of 63.35% and 24.53%, respectively, in the HEK293T dataset. In pairwise comparison, Amaranth elevated precision by 16% compared to Scallop2, and by 177% compared to StringTie2. In the mouse fibroblast dataset, Amaranth achieved 60.26% average precision, while Scallop2 and StringTie2 achieved 53.01% and 27.68% average precision, respectively, with the same sensitivity under control. In the mouse fibroblast dataset pairwise comparison, Amaranth outperformed Scallop2 in 309 (83.7%) cells. It elevated precision by 3.2% over Scallop2 and by 65% over StringTie2. Among all of the heuristic parameters, the accurate identification of the first exon contributed the most to the improvements of Amaranth (Note S2 and Tables S1-S8). The drop-out effect is one of the most prevalent problems in single-cell sequencing [40], and UMI reads can rule out many such cases. The UMI-guided pruning criteria combined contributed 31.3% relative improvement in precision with a sacrifice of reducing 8.1% matching transcripts (Fig. S4). We report the time and memory usage of all three different assembly tools (Note S4 and Figure S5). Amaranth uses more resources than Scallop2 and StringTie2. The whole Amaranth assembly pipeline on 192 human cells completes in 8 minutes on an 80-core machine.

**Figure 5.**
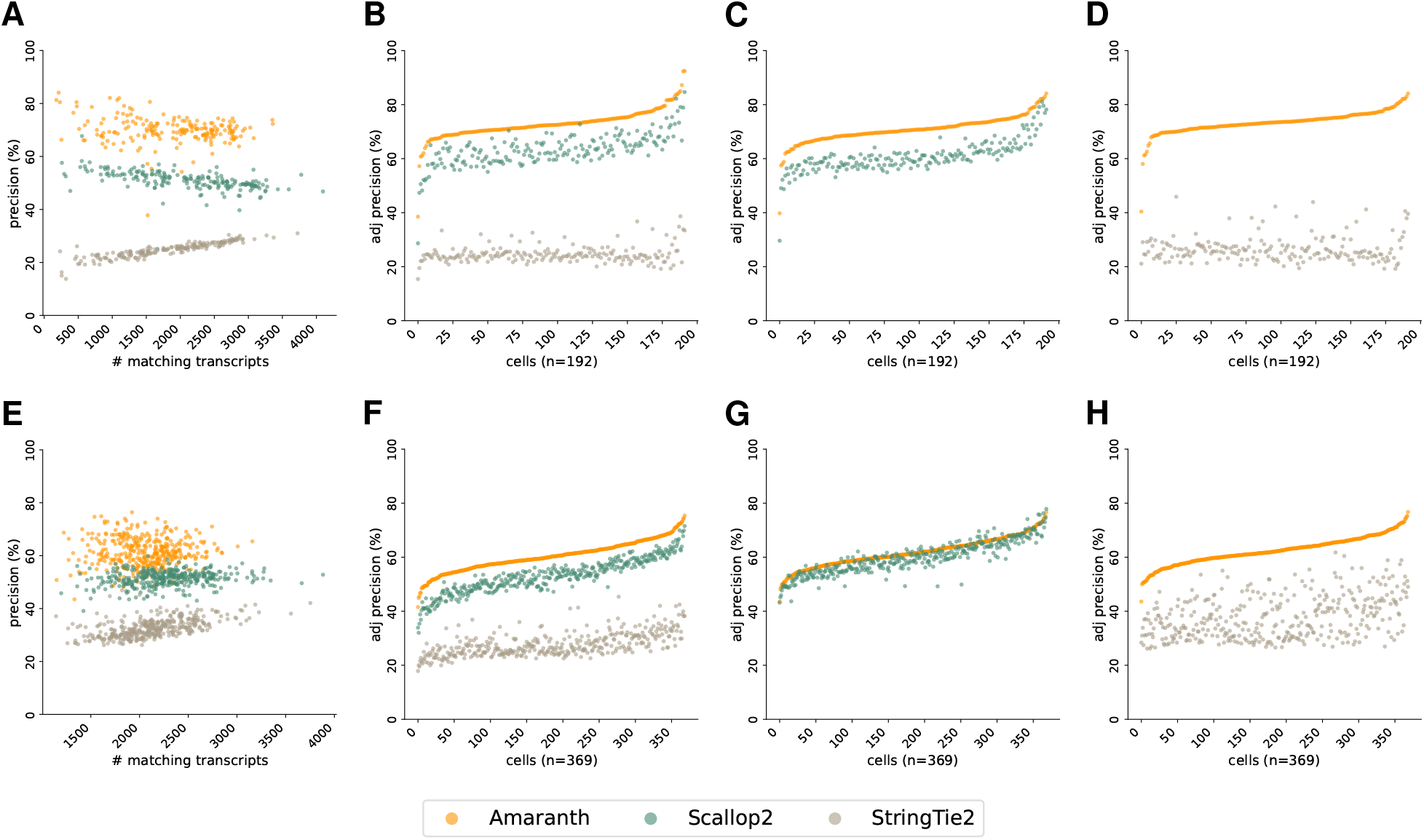
Comparison of assembly accuracy across all cells. **(A-D)** Comparison of assembly accuracy in HEK293T Smart-seq3 dataset. **(A)** Comparison of assembly precision and sensitivity across all cells. **(B)** Comparison of adjusted precision across all cells. Average adjusted precision: Amaranth 72.97%, Scallop2 63.35%, StringTie2 24.53%. **(C)** Pairwise comparison of adjusted precision of Amaranth and Scallop2. Average adjusted precision: Amaranth 70.75%, Scallop2 60.86%. Amaranth outperformed Scallop2 in 192 of 192 samples (100.0%). **(D)** Pairwise comparison of adjusted precision of Amaranth and StringTie2. Average adjusted precision: Amaranth 73.33%, StringTie2 26.48%. Amaranth outperformed StringTie2 in 192 of 192 samples (100.0%). **(E-H)** Comparison of assembly accuracy in mouse fibroblast Smart-seq3 dataset. **(E)** Comparison of assembly precision and sensitivity across all cells. **(F)** Comparison of adjusted precision across all cells. Average adjusted precision: Amaranth 60.26%, Scallop2 53.01%, StringTie2 27.68%. **(G)** Pairwise comparison of adjusted precision of Amaranth and Scallop2. Average adjusted precision: Amaranth 61.78%; Scallop2 59.86%. Amaranth outperformed Scallop2 in 309 of 369 samples (83.7%). **(H)** Pairwise comparison of adjusted precision of Amaranth and StringTie2. Average adjusted precision: Amaranth 62.43%, StringTie2 37.78%. Amaranth outperformed StringTie2 in 369 of 369 samples (100.0%). Each dot represents one individual cell assembled by an evaluated assembler. Cells in **(B-D)** and **(F-H)** are sorted in increasing order with respect to Amaranth’s adjusted precisions.

Those benchmark experiments and observations demonstrate the superiority and effectiveness of our novel discriminative model in the assembly of UMI reads and internal reads together.

### 4.3 Amaranth-meta improves meta-assembly of scRNA-seq

We compared Amaranth-meta with Aletsch [30] and PsiCLASS [34], two state-of-the-art meta-assemblers, that consistently outperform on both real data and simulation tests in previous studies [30, 31]. While different assemblers trade off differently with respect to recall and precision, respectively, with their default parameters, Amaranth-meta notably outputs many more matching transcripts than the other assemblers, regardless of datasets (Fig. 6A and E). Although PsiCLASS has the highest default precision, it has many fewer matching transcripts than Amaranth-meta. Both precision and recall of Aletsch are lower than those of Amaranth-meta. Pairwise comparison under controlled sensitivity shows that Amaranth-meta’s adjusted precision is higher than PsiCLASS in the human HEK293T dataset by relatively 2.2%, and much higher than PsiCLASS in the mouse fibroblast dataset by 36% (see Fig. 6D and H). Amaranth-meta outperforms Aletsch in almost all cells and outperforms PsiCLASS in most of the cells (78.1% of human HEK293T cells, and 99.7% mouse fibroblast cells).

**Figure 6.**
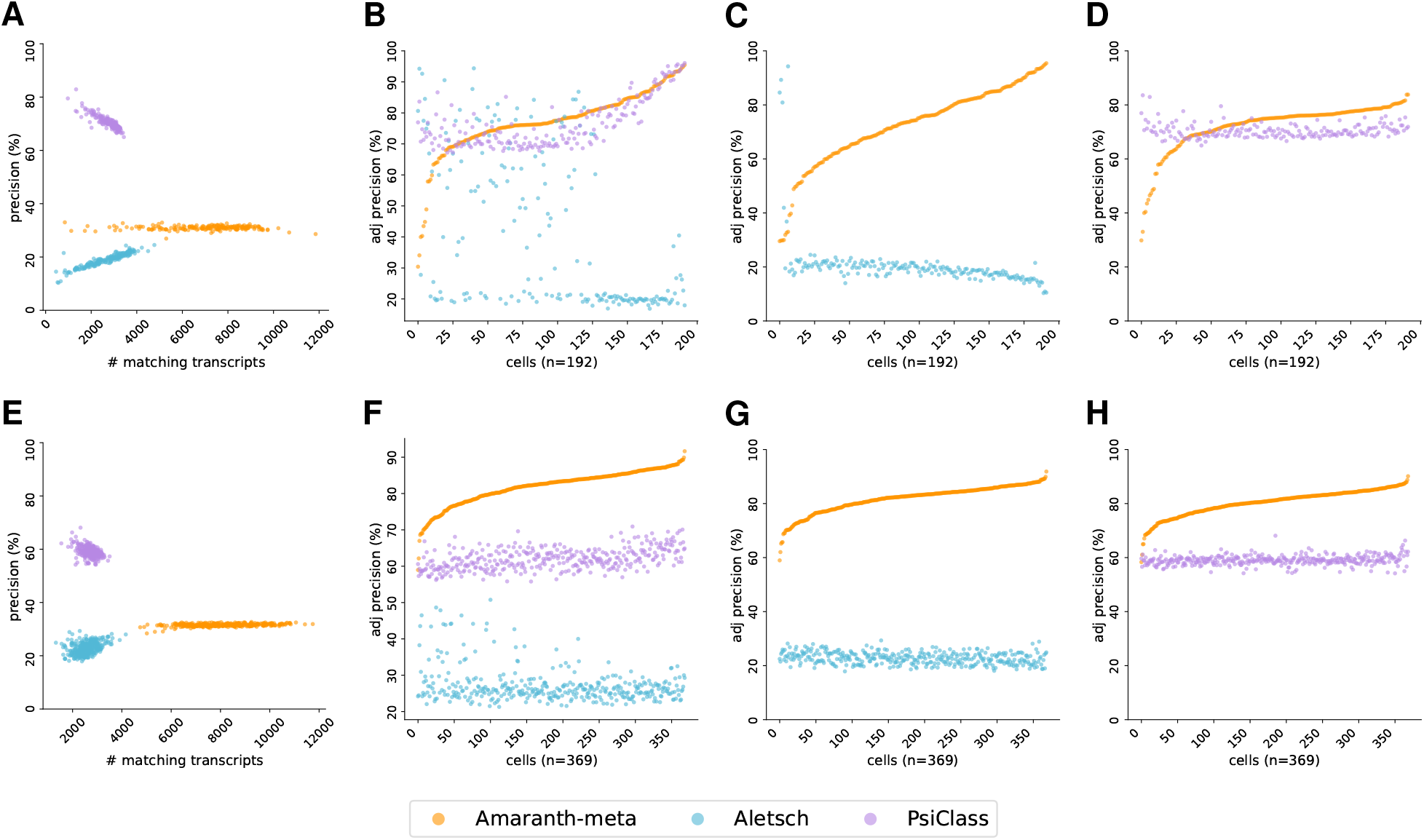
Comparison of meta-assembly accuracy. **(A-D)** Human HEK293T dataset. **(A)** Comparison of assembly precision and sensitivity across all cells. **(B)** Comparison of adjusted precision across all cells. Average adjusted precision: Amaranth-meta 77.06% precision, Aletsch 41.20%, PsiCLASS 76.55%. **(C)** Pairwise comparison of adjusted precisions of Amaranth-meta and Aletsch. Average adjusted precision: Amaranth-meta 72.34%, Aletsch 20.47%. Amaranth-meta outperformed Aletsch in 186 of 192 samples (96.9%). **(D)** Pairwise comparison of adjusted precisions of Amaranth-meta and PsiCLASS. Average adjusted precision: Amaranth-meta 72.05%, PsiCLASS 70.45%. Amaranth-meta outperformed PsiCLASS in 150 of 192 samples (78.1%). **(E-H)** Mouse fibroblast dataset. **(E)** Comparison of assembly precision and sensitivity across all cells. **(F)** Comparison of adjusted precision of all methods. Average adjusted precision: Amaranth-meta 81.86%, Aletsch 27.21%, PsiCLASS 62.02%. **(G)** Pairwise comparison of adjusted precisions of Amaranth-meta and Aletsch. Average adjusted precision: Amaranth-meta 81.70%, Aletsch 22.80%. Amaranth-meta outperformed Aletsch in 369 of 369 samples (100.0%). **(H)** Pairwise comparison of adjusted precision of Amaranth-meta and PsiCLASS. Average adjusted precision: Amaranth-meta 80.36%; PsiCLASS 59.05%. Amaranth-meta outperformed PsiCLASS in 368 of 369 samples (99.7%). Each dot represents one individual cell assembled by an evaluated assembler. Cells in **(B-D)** and **(F-H)** are sorted in increasing order with respect to Amaranth’s adjusted precisions.

## 5 Conclusions and Discussions

Single-cell RNA-seq has revolutionized transcriptome analysis, but full-length transcript reconstruction remains nevertheless challenging. While technologies like Smart-seq3 combine UMI-linked reads and internal reads to improve coverage, existing assembly tools fail to leverage their distinct biological and statistical properties. UMI reads exhibit 5’-end bias and high specificity, while internal reads provide broader coverage but introduce more noise. Hence, the distinct properties of reads in scRNA-seq assembly have yet to be fully exploited.

We present Amaranth, a novel scRNA-seq assembler that discriminatively integrates UMI and internal reads to achieve accurate, cell-specific transcript modeling. Amaranth’s key innovation lies in its multi-tiered computational framework that: (1) distinguishes and corrects read types while filtering PCR duplicates and contaminants, (2) constructs splice graphs while purging intronic contamination, and (3) leverages UMI read termini to accurately identify transcript start sites. By explicitly modeling the distinct properties of read types, Amaranth reduces false positives from non-specific internal reads while recovering transcripts that might be missed by approaches that rely solely on UMI reads.

Benchmarked on Smart-seq3 datasets from human HEK293T and mouse fibroblast cells, Amaranth and Amaranth-meta outperformed state-of-the-art tools, including StringTie2, Scallop2, Aletsch, and PsiCLASS, achieving superior precision by approximately 15%. Our results demonstrate that discriminative modeling of read types is crucial for accurate single-cell transcript assembly, enabling isoform-level resolution in scRNA-seq analysis. This advancement opens new possibilities for studying cell-type-specific splicing patterns and isoform regulation in heterogeneous cell populations.

While Amaranth demonstrates significant improvements in single-cell transcript assembly, it is currently optimized for Smart-seq series and similar protocols that generate both UMI reads and internal reads simultaneously. The performance on other scRNA-seq technologies would require specific adaptations and integration of extra data sources, for example, combining 10X genomics end-biased scRNA-seq data with transcriptome annotations. A promising future direction is the development of a hybrid assembly method that jointly leverages UMI-linked reads from single-cell protocols and internal reads from bulk RNA-seq datasets. Such an approach would allow exploiting the vast amount of existing bulk RNA-seq data alongside widely available 10x single-cell profiles, enabling large-scale and more comprehensive single-cell transcript assembly. The improvements achieved by Amaranth demonstrate the feasibility of such a hybrid assembly.

Amaranth achieved much improved precision on Smart-seq3 single-cell datasets. Although the UMI-guided pruning and intron removal procedure could inadvertently discard genuine intron-retention transcripts that carry biological functions or reflect cell-type-specific splicing patterns, decoupling assembly from intron retention identification reduces the already high complexity of transcript assembly and improves overall performance. Intron retention detection can instead be addressed downstream using dedicated alternative splicing analysis tools such as rMATS [29].

We acknowledge that though Amaranth-meta’s algorithm is straightforward and works well, it can benefit from a more delicate design of meta-information utilization. For example, constructing individual cell’s splice graphs with selective augmentation from others will produce more specific and precise transcripts than using a common splice graph for all cells, which maximizes the gain from other cells while limiting noise.

## Availability

Amaranth is implemented in C++ and is freely available as open-source software under the BSD-3-Clause license at https://github.com/Shao-Group/amaranth. Scripts, documentation, and data for reproducing the experiments in this manuscript are available at https://github.com/Shao-Group/amaranth-test.

## Acknowledgement

The authors thank Qimin Zhang for help in data collection and pre-processing. This work is supported by the US National Science Foundation (2145171 to M.S.) and by the US National Institutes of Health (R01HG011065 to M.S.).

## References

[1] Batzoglou, S., Pachter, L., Mesirov, J.P., Berger, B., Lander, E.S.: Human and Mouse Gene Structure: Comparative Analysis and Application to Exon Prediction. Genome Research 10(7), 950–958 (2000)

[2] Breschi, A., Gingeras, T.R., Guigó, R.: Comparative transcriptomics in human and mouse. Nature reviews. Genetics 18(7), 425–440 (2017)

[3] Broad Institute: Picard toolkit (2019), https://broadinstitute.github.io/picard/

[4] Chow, A., Lareau, C.A.: Concepts and new developments in droplet-based single-cell multiomics. Trends in biotechnology 42(11), 1379–1395 (2024)

[5] Dobin, A., Davis, C., Schlesinger, F., Drenkow, J., Zaleski, C., Jha, S., Batut, P., Chaisson, M., Gingeras, T.: STAR: ultrafast universal RNA-seq aligner. Bioinformatics 29(1), 15–21 (2013)

[6] Efremova, M., Vento-Tormo, M., Teichmann, S.A., Vento-Tormo, R.: CellPhoneDB: Inferring cell–cell communication from combined expression of multi-subunit ligand–receptor complexes. Nature Protocols 15(4), 1484–1506 (2020)

[7] Gupta, I., Collier, P.G., Haase, B., Mahfouz, A., Joglekar, A., Floyd, T., Koopmans, F., Barres, B., Smit, A.B., Sloan, S.A., et al.: Single-cell isoform RNA sequencing characterizes isoforms in thousands of cerebellar cells. Nature Biotechnology 36(12), 1197–1202 (2018)

[8] Hagemann-Jensen, M., Ziegenhain, C., Sandberg, R.: Scalable single-cell RNA sequencing from full transcripts with Smart-seq3xpress. Nat Biotechnol 40, 1452–1457 (2022)

[9] Hagemann-Jensen, M., Ziegenhain, C., Chen, P., Ramsköld, D., Hendriks, G.J., Larsson, A.J.M., Faridani, O.R., Sandberg, R.: Single-cell RNA counting at allele and isoform resolution using Smart-seq3. Nature Biotechnology 38, 708–714 (2020)

[10] Hahaut, V., Pavlinic, D., Carbone, W., Schuierer, S., Balmer, P., Quinodoz, M., Renner, M., Roma, G., Cowan, C.S., Picelli, S.: Fast and highly sensitive full-length single-cell rna sequencing using flash-seq. Nature Biotechnology 40(10), 1447–1451 (2022)

[11] Hayashi, T., Ozaki, H., Sasagawa, Y., Umeda, M., Danno, H., Nikaido, I.: Single-cell full-length total rna sequencing uncovers dynamics of recursive splicing and enhancer rnas. Nature Communications 9(1), 619 (2018)

[12] Kim, D., Langmead, B., Salzberg, S.: HISAT: a fast spliced aligner with low memory requirements. Nat. Methods 12(4), 357–360 (2015)

[13] Kim, D., Paggi, J.M., Park, C., Bennett, C., Salzberg, S.L.: Graph-based genome alignment and genotyping with HISAT2 and HISAT-genotype. Nature Biotechnology 37(8), 907–915 (2019)

[14] Kovaka, S., Zimin, A.V., Pertea, G.M., Razaghi, R., Salzberg, S.L., Pertea, M.: Transcriptome assembly from long-read RNA-seq alignments with StringTie2. Genome Biology 20, 278 (2019)

[15] La Manno, G., Soldatov, R., Zeisel, A., Braun, E., Hochgerner, H., Petukhov, V., Lidschreiber, K., Kastriti, M.E., Lönnerberg, P., Furlan, A., Fan, J., Borm, L.E., Liu, Z., van Bruggen, D., Guo, J., He, X., Barker, R., Sundström, E., Castelo-Branco, G., Cramer, P., Adameyko, I., Linnarsson, S., Kharchenko, P.V.: RNA velocity of single cells. Nature 560(7719), 494–498 (2018)

[16] Li, H.: Minimap2: pairwise alignment for nucleotide sequences. Bioinformatics 34(18), 3094–3100 (2018)

[17] Liu, J., Yu, T., Jiang, T., Li, G.: TransComb: genome-guided transcriptome assembly via combing junctions in splicing graphs. Genome Biol. 17(1), 213 (2016)

[18] Liu, J., Liu, X., Ren, X., Li, G.: scRNAss: a single-cell RNA-seq assembler via imputing dropouts and combing junctions. Bioinformatics 35(21), 4264–4271 (2019)

[19] Macosko, E.Z., Basu, A., Satija, R., Nemesh, J., Shekhar, K., Goldman, M., Tirosh, I., Bialas, A.R., Kamitaki, N., Martersteck, E.M., Trombetta, J.J., Weitz, D.A., Sanes, J.R., Shalek, A.K., Regev, A., McCarroll, S.A.: Highly Parallel Genome-wide Expression Profiling of Individual Cells Using Nanoliter Droplets. Cell 161(5), 1202–1214 (2015)

[20] Nip, K.M., Chiu, R., Yang, C., Chu, J., Mohamadi, H., Warren, R.L., Birol, I.: RNA-Bloom enables reference-free and reference-guided sequence assembly for single-cell transcriptomes. Genome Research 30, 1191–1200 (2020)

[21] Parekh, S., Ziegenhain, C., Vieth, B., Enard, W., Hellmann, I.: zUMIs - a fast and flexible pipeline to process RNA sequencing data with UMIs. GigaScience 7(6), giy059 (Jun 2018)

[22] Pertea, G., Pertea, M.: Gff utilities: Gffread and gffcompare. F1000Research 9 (2020)

[23] Picelli, S., Faridani, O.R., Åsa K Björklund, Winberg, G., Sagasser, S., Sandberg, R.: Full-length rna-seq from single cells using Smart-seq2. Nature Protocols 9, 171–181 (2014)

[24] Rodriques, S.G., Stickels, R.R., Goeva, A., Martin, C.A., Murray, E., Vanderburg, C.R., Welch, J., Chen, L.M., Chen, F., Macosko, E.Z.: Slide-seq: A scalable technology for measuring genome-wide expression at high spatial resolution. Science 363(6434), 1463–1467 (2019)

[25] Rosenberg, A.B., Roco, C.M., Muscat, R.A., Kuchina, A., Sample, P., Yao, Z., Graybuck, L.T., Peeler, D.J., Mukherjee, S., Chen, W., Pun, S.H., Sellers, D.L., Tasic, B., Seelig, G.: Single-cell profiling of the developing mouse brain and spinal cord with split-pool barcoding. Science 360(6385), 176–182 (2018)

[26] Roy, S.W., Fedorov, A., Gilbert, W.: Large-scale comparison of intron positions in mammalian genes shows intron loss but no gain. Proceedings of the National Academy of Sciences 100(12), 7158–7162 (2003)

[27] Salmen, F., De Jonghe, J., Kaminski, T.S., Alemany, A., Parada, G.E., Verity-Legg, J., Yanagida, A., Kohler, T.N., Battich, N., van den Brekel, F., Ellermann, A.L., Arias, A.M., Nichols, J., Hemberg, M., Hollfelder, F., van Oudenaarden, A.: High-throughput total RNA sequencing in single cells using VASA-seq. Nature Biotechnology pp. 1–14 (2022)

[28] Shao, M., Kingsford, C.: Accurate assembly of transcripts through phase-preserving graph decomposition. Nature Biotechnology 35(12), 1167–1169 (2017)

[29] Shen, S., Park, J.W., Lu, Z.x., Lin, L., Henry, M.D., Wu, Y.N., Zhou, Q., Xing, Y.: rMATS: Robust and flexible detection of differential alternative splicing from replicate RNA-Seq data. Proceedings of the National Academy of Sciences 111(51), E5593–E5601 (2014)

[30] Shi, Q., Zhang, Q., Shao, M.: Accurate assembly of multiple RNA-seq samples with Aletsch. Bioinformatics 40(Supplement_1), i307–i317 (2024)

[31] Shi, Q., Zhang, Q., Shao, M.: Transcriptome Assembly at Single-Cell Resolution with Beaver (2024)

[32] Shi, Z.X., Chen, Z.C., Zhong, J.Y., Hu, K.H., Zheng, Y.F., Chen, Y., Xie, S.Q., Bo, X.C., Luo, F., Tang, C., et al.: High-throughput and high-accuracy single-cell rna isoform analysis using pacbio circular consensus sequencing. Nature Communications 14(1), 2631 (2023)

[33] Song, L., Sabunciyan, S., Florea, L.: CLASS2: accurate and efficient splice variant annotation from RNA-seq reads. Nucleic Acids Res 44(10), e98 (2016)

[34] Song, L., Sabunciyan, S., Yang, G., Florea, L.: A multi-sample approach increases the accuracy of transcript assembly. Nature communications 10(1), 5000 (2019)

[35] Ståhl, P.L., Salmén, F., Vickovic, S., Lundmark, A., Navarro, J.F., Magnusson, J., Giacomello, S., Asp, M., Westholm, J.O., Huss, M., Mollbrink, A., Linnarsson, S., Codeluppi, S., Borg, Å., Pontén, F., Costea, P.I., Sahlén, P., Mulder, J., Bergmann, O., Lundeberg, J., Frisén, J.: Visualization and analysis of gene expression in tissue sections by spatial transcriptomics. Science 353(6294), 78–82 (2016)

[36] Trapnell, C., Williams, B., Pertea, G., Mortazavi, A., Kwan, G., Van Baren, M., Salzberg, S., Wold, B., Pachter, L.: Transcript assembly and quantification by RNA-Seq reveals unannotated transcripts and isoform switching during cell differentiation. Nat. Biotechnol. 28(5), 511–515 (2010)

[37] Trapnell, C.: Defining cell types and states with single-cell genomics. Genome Research 25(10), 1491–1498 (2015)

[38] Wang, L., Wang, S., Li, W.: RSeQC: Quality control of RNA-seq experiments. Bioinformatics 28(16), 2184–2185 (2012)

[39] Wang, X., He, Y., Zhang, Q., Ren, X., Zhang, Z.: Direct comparative analyses of 10x genomics chromium and smart-seq2. Genomics, Proteomics and Bioinformatics 19(2), 253–266 (2021)

[40] Westoby, J., Artemov, P., Hemberg, M., Ferguson-Smith, A.: Obstacles to detecting isoforms using full-length scRNA-seq data. Genome Biology 21(1), 74 (2020)

[41] Yu, T., Zhao, X., Li, G.: Transmeta simultaneously assembles multisample rna-seq reads. Genome Research 32(7), 1398–1407 (2022)

[42] Zhang, Q., Shao, M.: Transcript assembly and annotations: Bias and adjustment. PLOS Computational Biology 19(12), e1011734 (2023)

[43] Zhang, Q., Shi, Q., Shao, M.: Accurate assembly of multi-end rna-seq data with scallop2. Nature Computational Science 2(3), 148–152 (2022)

[44] Zheng, G.X.Y., Terry, J.M., Belgrader, P., Ryvkin, P., Bent, Z.W., Wilson, R., Ziraldo, S.B., Wheeler, T.D., McDermott, G.P., Zhu, J., Gregory, M.T., Shuga, J., Montesclaros, L., Underwood, J.G., Masquelier, D.A., Nishimura, S.Y., Schnall-Levin, M., Wyatt, P.W., Hindson, C.M., Bharadwaj, R., Wong, A., Ness, K.D., Beppu, L.W., Deeg, H.J., McFarland, C., Loeb, K.R., Valente, W.J., Ericson, N.G., Stevens, E.A., Radich, J.P., Mikkelsen, T.S., Hindson, B.J., Bielas, J.H.: Massively parallel digital transcriptional profiling of single cells. Nature Communications 8(1), 14049 (2017)

