## Supplemental materials for "Amaranth: Enhanced Single-Cell Transcript Assembly via Discriminative Modeling of UMI Reads and Internal Reads"

### S1 Parameters of Amaranth

- `--min_umi_reads_bundle` (*int*, default: 1): Bundle (gene locus) with fewer UMI reads than this threshold will be ignored.
- `--min_umi_ratio_bundle` (*float*, default: 0.0): Bundle (gene locus) with lower UMI reads ratio than this threshold will be ignored.
- `--both_umi_support` (*flag*, default: off): If set, a bundle needs to satisfy both UMI support thresholds (`--min_umi_reads_bundle` and `--min_umi_ratio_bundle`). Otherwise, satisfying either is sufficient.
- `--min_umi_reads_start_exon` (*int*, default: 1): Minimum number of UMI reads supporting the first exon in a valid transcript.
- `--remove-retained-intron` (*flag*, default: on): Remove retained introns. To disable this option, use `--no-remove-retained-intron`.
- `--no-remove-retained-intron` (*flag*, default: off): Do not remove retained introns.
- `--remove-pcr-duplicates` (*int*, default: 1): Option `0`: do not remove; Or option `1`: remove PCR duplicates with identical alignment coordinates and CIGAR string.
- `--max-ir-part-ratio-v` (*float*, default: 0.5): The ratio threshold of retained node to skip edge for partial introns. If greater than threshold, consider true transcript.
- `--max-ir-part-ratio-e` (*float*, default: 0.5): The ratio threshold of retained node's edge to skip edge for partial introns. If greater than threshold, consider true transcript.
- `--max-ir-full-ratio-v` (*float*, default: 1.0): The ratio threshold of retained node to skip edge for full introns. If greater than threshold, consider true transcript.
- `--max-ir-full-ratio-e` (*float*, default: 0.5): The ratio threshold of retained node's edge to skip edge for full introns. If greater than threshold, consider true transcript.
- `--max-ir-full-ratio-i` (*float*, default: 10.0): The ratio threshold of retained node to its own edge for full introns. If greater than threshold, consider true retention.
- `--max-ir-umi-support-full` (*int*, default: 3): The minimum number of UMI reads to support a partial exon rather than a full intron retention. If lower than the threshold, consider true retention.
- `--max-ir-umi-support-part` (*int*, default: 5): The minimum number of UMI reads to support a partial exon rather than partial intron retention. If lower than the threshold, consider true retention.
- `--min-cb-ratio` (*double*, default: 0.3): For meta-assembly only, minimum ratio of exons in a transcript that is supported by a cell's barcode (CB) to assign this transcript to this cell.

### S2 Ablation experiments of heuristic parameters

Ablation experiments were performed for each of the tunable parameters described in the Amaranth algorithm. Eight parameters were examined; per-parameter ablation results on human HEK293T cells (192 cells) are reported in Tables S1–S8. Each table varies one parameter while holding the rest at their most permissive values (i.e., 0) when applicable, so that the performance change can be mostly attributed to the variable under investigation. When evaluating `--min_umi_ratio_bundle` and `--min_umi_reads_bundle`, the parameter `--both-umi-support` is set to true, because in the default (either-sufficient) mode, a bundle satisfying any single criterion is retained, making it impossible to isolate the effect of the parameter under test. The asterisk (*) marks the value used in default settings and in the main experiments. Precision (%) and number of matching transcripts are reported as mean ± standard deviation across cells. It is noteworthy that the effects of different parameters are neither independent nor linear, making it difficult to evaluate the efficacy of each in isolation. The drop-out event is one of the most prevalent problems in single-cell sequencing, while the good performance of `--min_umi_reads_start_exon` indicates UMI reads are strong indicators for full-length transcripts, and distinguishing UMI and internal reads improves assembly quality. For reference, Amaranth has 68.96% precision and 1902 matching transcripts under default settings.

### S3 Ablation experiment of UMI-guided pruning

Ablation experiments were performed on 192 human HEK293T cells to evaluate UMI-guided pruning effects. Specifically, Amaranth was run with all default parameters as UMI-guided pruning mode, and run with all permissive-value parameters, that were used in Section S2, as non-pruning mode. Fig. S4 shows the precisions and number of matching transcripts of UMI-guided pruning and non-pruning. In conclusion, the UMI-guided pruning criteria combined contributed 31.3% relative improvement in precision (from 52.49 percentage points to 68.96 percentage points) with a sacrifice of reducing 8.1% matching transcripts (from 2072 to 1902), indicating that UMI support filtering effectively removes false-positive assemblies while retaining the majority of true transcripts.

### S4 Time and memory usage of assembly tools

Time and memory usage of three assembly tools were reported in Fig. S5 using Nextflow's trace information (`realtime` and `peak_rss`). Amaranth uses the most resources (median runtime 120 s and median peak memory 46.5 MB per cell), but remains at an affordable level. Assembling all those cells on an 80-core HPC using Amaranth takes 8 min and less than 10 GB peak memory.

### Supplementary Figures

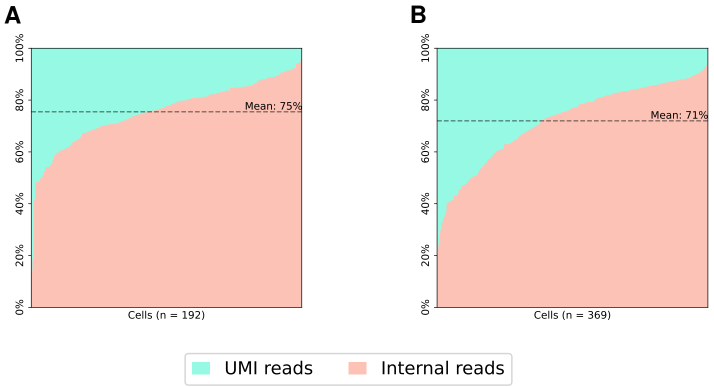


Fig. S1: **Proportion of internal and UMI reads in human HEK293T cells and mouse fibroblast cells.** Cells are sorted in ascending order of the proportion of internal reads. **(A)** Proportion of reads in human HEK293T cells (192 cells). On average, 75% of reads in a HEK293T cell are internal, while only 25% of reads are UMI reads. **(B)** Proportion of reads in mouse fibroblast cells (369 cells). On average, 71% of reads in a mouse fibroblast cell are internal, while only 29% of reads are UMI reads.

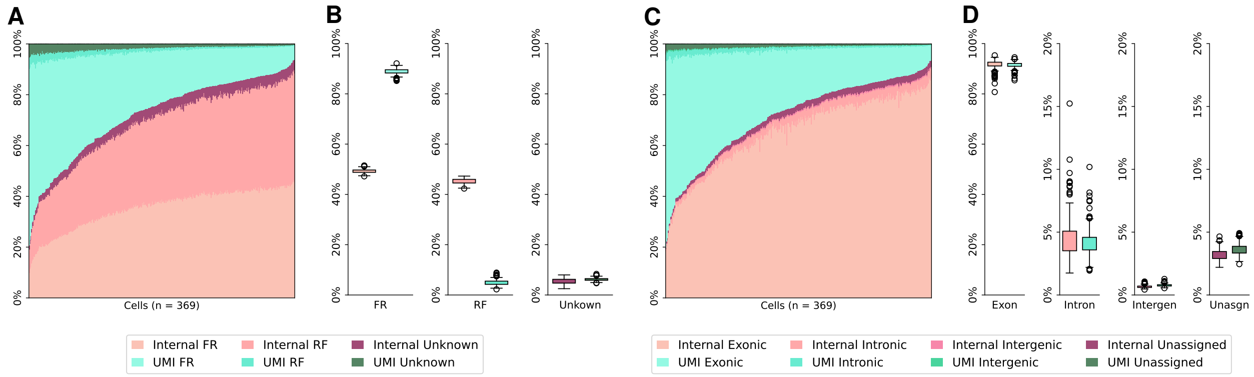


Fig. S2: **Read strandness and genome feature distributions of UMI reads and internal reads in mouse fibroblast cells. (A)** Strandness of reads in each mouse fibroblast cell. Cells are sorted in ascending order of internal reads proportion. **(B)** Breakdown of read strandness by read type. Internal reads: 49.24% FR-strandness, 45.26% RF-strandness, 5.50% unknown. UMI reads: 88.89% FR-strandness, 4.91% RF-strandness, 6.20% unknown. **(C)** Genome features of read tags in each mouse fibroblast cell. Cells are sorted in ascending order of internal reads proportion. **(D)** Breakdown of genome feature distributions by read type in mouse fibroblast cells. Internal read tags: 91.69% exonic, 4.45% intronic, 0.66% intergenic, 3.20% unassigned. UMI read tags: 91.45% exonic, 4.15% intronic, 0.77% intergenic, 3.63% unassigned.

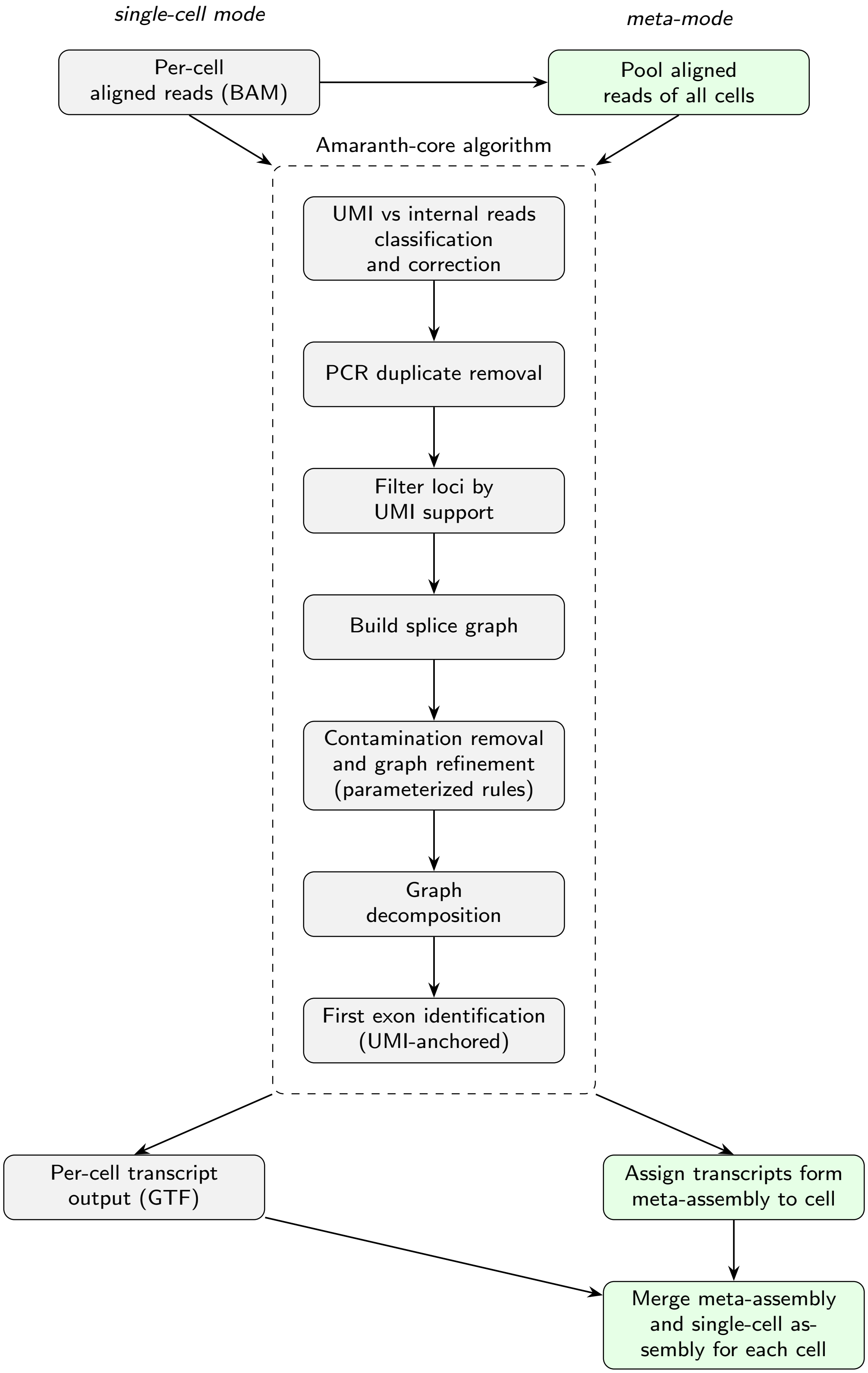


Fig. S3: **Amaranth workflow.** This figure illustrates the outline of Amaranth's single-cell mode (left) and meta-assembly mode (right) with the shared core assembly algorithm.

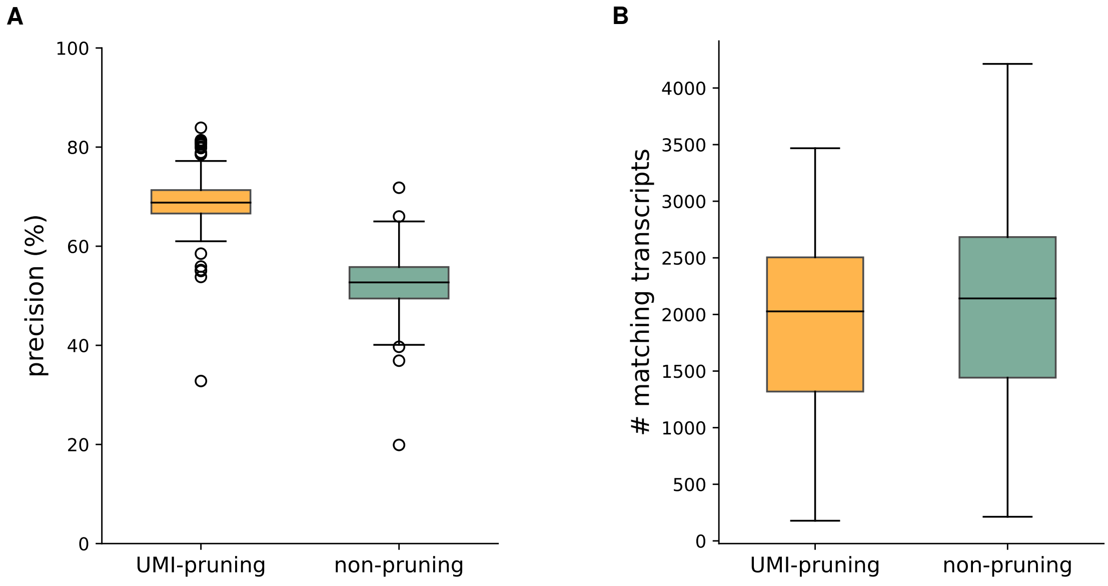


Fig. S4: **Effect of UMI-guided pruning on assembly performance in the HEK293T single-cell dataset (n = 192 cells)**. With UMI-pruning, mean precision is 68.96% with 1902 matching transcripts. Without UMI-pruning, mean precision is 52.49% with 2072 matching transcripts.

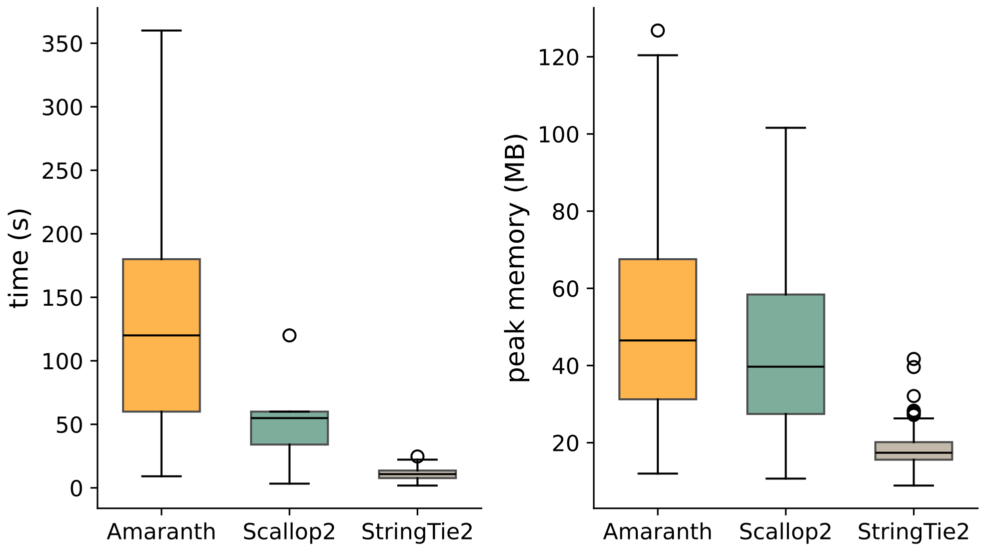


Fig. S5: **Time and memory usage of assembly tools on each of 192 cells in HEK293T dataset**. Box plots show the distribution of wall-clock time and peak memory for Amaranth, Scallop2 and StringTie2, each with n=192 cells. Amaranth had a median runtime of 120.0 s and median peak memory of 46.5 MB. Scallop2 had a median runtime of 54.9 s and median peak memory of 39.7 MB. StringTie2 had a median runtime of 10.8 s and median peak memory of 17.4 MB.

### Supplementary Tables

| Value | Precision (%) | # matching transcripts |
|---|---|---|
| 0 | $52.49 \pm 5.65$ | $2071.9 \pm 826.7$ |
| 0.1 | $53.31 \pm 5.38$ | $2069.1 \pm 825.0$ |
| $0.5^*$ | $54.45 \pm 4.96$ | $2059.2 \pm 820.2$ |
| 1.0 | $54.86 \pm 4.80$ | $2054.3 \pm 818.3$ |
| 2.0 | $55.12 \pm 4.69$ | $2051.0 \pm 816.4$ |
| 5.0 | $55.29 \pm 4.60$ | $2044.9 \pm 812.6$ |

Table S1: Ablation of `-max-ir-full-ratio-e`: ratio threshold of retained node's edge to skip edge for full introns.

| Value | Precision (%) | # matching transcripts |
|---|---|---|
| 0 | $55.27 \pm 4.53$ | $2029.4 \pm 803.9$ |
| 2.0 | $52.49 \pm 5.65$ | $2071.9 \pm 826.7$ |
| 5.0 | $52.49 \pm 5.65$ | $2071.9 \pm 826.7$ |
| $10.0^*$ | $52.49 \pm 5.65$ | $2071.9 \pm 826.7$ |
| 20.0 | $52.49 \pm 5.65$ | $2071.9 \pm 826.7$ |

Table S2: Ablation of `-max-ir-full-ratio-i`: ratio threshold of retained node to its own edge for full introns.

| Value | Precision (%) | # matching transcripts |
|---|---|---|
| 0 | $52.49 \pm 5.65$ | $2071.9 \pm 826.7$ |
| 0.5 | $54.45 \pm 4.96$ | $2059.2 \pm 820.2$ |
| $1.0^*$ | $54.86 \pm 4.80$ | $2054.3 \pm 818.3$ |
| 2.0 | $55.12 \pm 4.69$ | $2051.0 \pm 816.4$ |
| 5.0 | $55.29 \pm 4.60$ | $2044.9 \pm 812.6$ |
| 10.0 | $55.33 \pm 4.56$ | $2039.5 \pm 809.5$ |

Table S3: Ablation of `-max-ir-full-ratio-v`: ratio threshold of retained node to skip edge for full introns.

| Value | Precision (%) | # matching transcripts |
|---|---|---|
| 0 | $52.49 \pm 5.65$ | $2071.9 \pm 826.7$ |
| 0.1 | $53.09 \pm 5.45$ | $2070.5 \pm 826.0$ |
| $0.5^*$ | $54.69 \pm 4.89$ | $2064.7 \pm 822.1$ |
| 1.0 | $56.76 \pm 4.14$ | $2003.4 \pm 788.1$ |
| 2.0 | $56.76 \pm 4.14$ | $2003.4 \pm 788.1$ |
| 5.0 | $56.76 \pm 4.14$ | $2003.4 \pm 788.1$ |

Table S4: Ablation of `-max-ir-part-ratio-e`: ratio threshold of retained node's edge to skip edge for partial introns.

| Value | Precision (%) | # matching transcripts |
|---|---|---|
| 0 | $52.49 \pm 5.65$ | $2071.9 \pm 826.7$ |
| 0.1 | $53.09 \pm 5.46$ | $2070.1 \pm 825.6$ |
| $0.5^*$ | $54.71 \pm 4.89$ | $2062.0 \pm 820.4$ |
| 1.0 | $56.77 \pm 4.16$ | $2021.4 \pm 796.3$ |
| 2.0 | $56.77 \pm 4.15$ | $2004.8 \pm 789.4$ |
| 5.0 | $56.77 \pm 4.14$ | $2003.5 \pm 788.2$ |

Table S5: Ablation of `-max-ir-part-ratio-v`: ratio threshold of retained node to skip edge for partial introns.

| Value | Precision (%) | # matching transcripts |
|---|---|---|
| 0 | $52.49 \pm 5.66$ | $2072.1 \pm 826.7$ |
| $1^*$ | $63.10 \pm 6.42$ | $1933.7 \pm 754.0$ |
| 2 | $69.52 \pm 7.01$ | $1287.5 \pm 505.3$ |
| 5 | $74.95 \pm 8.99$ | $597.6 \pm 254.9$ |

Table S6: Ablation of `-min_umi_reads_start_exon`: minimum number of UMI reads supporting the first exon in a valid transcript.

| Value | Precision (%) | # matching transcripts |
|---|---|---|
| $0.0^*$ | $52.49 \pm 5.67$ | $2071.8 \pm 826.6$ |
| 0.01 | $57.25 \pm 6.53$ | $2008.5 \pm 795.3$ |
| 0.05 | $57.77 \pm 6.23$ | $1927.3 \pm 735.0$ |
| 0.1 | $58.76 \pm 5.84$ | $1738.7 \pm 655.7$ |

Table S7: Ablation of `-min_umi_ratio_bundle`: minimum UMI reads ratio for a bundle (gene locus) to be retained.

| Value | Precision (%) | # matching transcripts |
|---|---|---|
| 0 | $52.49 \pm 5.67$ | $2071.8 \pm 826.6$ |
| $1^*$ | $57.22 \pm 6.56$ | $2010.0 \pm 797.5$ |
| 2 | $57.24 \pm 6.56$ | $2010.1 \pm 797.6$ |
| 5 | $62.71 \pm 8.41$ | $1760.7 \pm 722.0$ |

Table S8: Ablation of `-min_umi_reads_bundle`: minimum UMI reads for a bundle (gene locus) to be retained.
